# Females prefer cooperative males even when cooperative behavior is unobserved: evidence from the mound-building mouse, *Mus spicilegus*

**DOI:** 10.1101/197988

**Authors:** Arnaud Tognetti, Michel Raymond, Guila Ganem, Charlotte Faurie

## Abstract

Theoretical and empirical studies in humans suggest that cooperative behaviors may act as signals during mate choice. However, cooperation is not always observable by potential partners before mate choice. To address whether cooperative phenotypes are preferred based on cues different from cooperative behaviors *per se,* we designed an experimental paradigm using wild-born mound-building mice (*Mus spicilegus*), a species with biparental care. In this species, females cannot observe male cooperative behaviors: mate choice occurs in the spring, whereas mounds are cooperatively built in the fall. We first assessed the variation in mound building investment and identified males exhibiting high and low amounts of cooperation. Second, we presented these males to females during two-way choice tests. As offspring survival relies on mound protection, we hypothesized that mound building could be a form of paternal care and assessed whether cooperative males were more involved in offspring attendance using pup-retrieval experiments. Our results indicate that females were more attracted to highly cooperative males over less cooperative, even when they did not observe them build. This finding suggests that female mate choice is influenced either by cues of cooperativeness different than cooperative behaviors or by preferences for traits associated with cooperativeness. Moreover, male offspring attendance was negatively correlated with cooperativeness, suggesting the potential existence of two alternative paternal strategies in offspring care (mound building versus offspring attendance). Overall, our findings support the existence of preference for cooperative phenotypes in a non-human species and suggest that sexual selection might be involved in the evolution of cooperation.

## Introduction

Several theoretical studies have highlighted the potential role of the direct social and reproductive benefits of cooperation (Zahavi 1995; Putland 2001; McNamara et al. 2008; Barclay 2011; Wolf and Krause 2014). Cooperative behaviors were shown to vary among individuals (Heinsohn and Legge 1999; Clutton-Brock et al. 2002; Russell et al. 2003) and to influence mate choice, at least in humans. For example, both men and women seem to prefer cooperative mates (Farrelly et al. 2007; Phillips et al. 2008; Barclay 2010; Moore et al. 2013; Oda et al. 2013; Tognetti et al. 2014; Arnocky et al. 2016), with women being more sensitive to cooperativeness than men when choosing a sexual partner (Phillips et al. 2008). Cooperative individuals have also been shown to be rated as physically more attractive (Farrelly et al. 2007) and report more sexual partners (Arnocky et al. 2016). Moreover, men use cooperative behaviors as a signaling strategy: in the presence of women, they are more cooperative (Tognetti et al. 2012; Van Vugt and Iredale 2013), and they compete by increasing their cooperativeness in response to displays from competitors (Van Vugt and Iredale 2013; Raihani and Smith 2015; Tognetti et al. 2016). So far, in non-human species, no experimental evidence supports the potential role of mate choice in the evolution of cooperation (e.g: McDonald, Kazem, et al. 2008; McDonald, te Marvelde, et al. 2008; Nomano et al. 2013). However, when cooperative behaviors of potential mates cannot be directly observed by the chooser, signals or cues of cooperativeness may be selected for during mate choice. Evidence for traits that reliably advertise the propensity to cooperate does exist. In humans, morphological facial traits, such as facial width-to-height ratio, advertise cooperativeness and trustworthiness (Stirrat and Perrett 2010; Stirrat and Perrett 2012; Tognetti et al. 2013). A genuine smile also seems to be a trait used to estimate cooperativeness (Oda et al. 2009; Centorrino et al. 2015). In the barn owl (*Tyto alba*), the propensity to cooperate has recently been linked to melanin-based plumage (Roulin et al. 2012). Moreover, the association of cooperativeness with other traits that are sexually selected could also indirectly select for cooperative behavior. Indeed, behavioral traits often form a suite of correlated traits so-called behavioral syndrome (Sih, A.M. Bell, et al. 2004; Sih, A. Bell, et al. 2004; Réale et al. 2010). For example, aggressiveness is associated positively with exploratory behavior and boldness in a number of species (Sih, A.M. Bell, et al. 2004; Sih, A. Bell, et al. 2004; Groothuis and Carere 2005; Boon et al. 2008; Réale et al. 2009) and negatively associated with parental care (Mutzel et al. 2013). These behavioral correlations can result from natural and sexual selection that favor particular trait combinations (Sih, A. Bell, et al. 2004; Réale et al. 2010; Schuett et al. 2010; Pruitt et al. 2011; Schuett et al. 2011; Kortet et al. 2012). For example, by preferring exploratory males, exploratory zebra finch females (*Taeniopygia guttata*) simultaneously select for more aggressive males due to an association between those traits (Schuett et al. 2011). The same process might select for cooperative tendencies in social species. For example, some studies suggest that cooperativeness is positively associated with paternal care and negatively associated with aggressiveness (Bergmüller et al. 2010; Réale et al. 2010). Hence, females that prefer less aggressive males or males engaging in more parental care would be indirectly selecting for cooperative males. In this way, cooperativeness could be selected for without being a mate choice criterion if another trait that is associated with it is sexually preferred.

The existence of cues that reliably advertise an individual's cooperativeness, and the possibility for cooperativeness to be indirectly selected for through its association with sexually selected traits, both stress the need to pursue investigations on the potential role of mate choice and possibly sexual selection in the evolution of cooperation. The main aim of our study is to test whether male cooperative phenotypes are preferred by females in a mouse, *Mus spicilegus*, where mate choice occurs before mound building and, hence, before the potential display of male cooperative behaviors.

*Mus spicilegus* is a socially monogamous species, endemic to southeastern Europe, and it possesses a cooperative mound building behavior that is unique among mice. After a period of reproduction from spring to late summer, during which adult breeding pairs produce a few litters in burrows of simple design (Sokolov et al. 1998), several breeding pairs gather in early autumn to build a common overwintering structure in which they collectively deposit their last litter in a nest chamber located under the mound (Muntyanu 1990; Garza et al. 1997; Poteaux et al. 2008; Canady et al. 2009). The mounds are built by the accumulation of plant materials covered with earth and are up to four meters in diameter and typically 0.5 meters high when freshly built (Muntyanu 1990; Sokolov et al. 1998). Adults have rarely been found in mounds during the winter, which suggests that their lifespans do not exceed 10 months (Muntyanu 1990; Poteaux et al. 2008; Canady et al. 2009). Only their offspring, overwintering under the mound, will thus establish the next generation the following year (Muntyanu 1990; Poteaux et al. 2008; Canady et al. 2009). These overwintering juveniles utilize the mound from autumn to spring, and do not reproduce within it. At the beginning of the breeding season, they leave the mound and form breeding pairs. A genetic approach revealed that among the breeding pairs that gather and deposit their offspring in a given mound, some mothers were genetically related whereas all fathers were unrelated (Garza et al. 1997; Poteaux et al. 2008). Although kin selection might be involved in female communal nesting, such a mechanism does not seem sufficient to explain cooperative mound building by unrelated males. We hence examine whether direct fitness benefits, such as enhanced mating success, may also be involved in the evolution of cooperative building.

Previous studies have found that, in captivity, investment in mound building varies between individuals (Simeonovska-Nikolova and Mehmed 2009; Serra et al. 2012) and that mound size is highly variable in natural conditions (Holzl et al. 2009). Since mound size is positively correlated with water insulation and the soil temperature inside the mound, it is also expected to influence the probability of offspring survival during the winter (Szenczi et al. 2011; Szenczi et al. 2012). It is therefore likely that females would benefit from choosing a partner that invests greatly in mound building. Nevertheless, knowledge about the behavior of this species in the wild is scarce, and both mate preference and cooperativeness-signaling mechanisms remain unknown. Notably, females select a partner in the spring, before seeing male investment in mound building that occurs a few months later (Muntyanu 1990; Sokolov et al. 1998). Hence, the ability of females to detect male cooperativeness through other signals/cues of cooperativeness should be positively selected for. Alternatively, females could indirectly select for male cooperation through their preferences for other traits associated with the male propensity to cooperate.

Cooperative investment in mound building could be seen as a form of parental care. Behavioral experiments in *Mus spicilegus* revealed that males invest in offspring attendance and that highly attendant fathers increase their reproductive success, as well as the reproductive success of their mate (Patris and Baudoin 2000; Feron and Gouat 2007). Hence, another aim of this study is to test whether males that invest highly in mound building also attend more to their offspring, or if these two forms of paternal investment are uncoupled.

We designed our experiment to test whether females preferred males who cooperated in mound building when cooperation was observed versus unobserved, and whether male cooperativeness in mound building was positively associated with paternal attendance to offspring. We assessed male spontaneous cooperativeness exhibited during mound building in captivity with wild mice captured in agricultural fields in Bulgaria. Then, we used two-way choice tests to assess females' preference for males with high values of investment in mound building versus males with low values. Finally, we evaluated the post-mating paternal care exhibited by males with varied levels of cooperativeness using pup retrieval experiments: we recorded the number of pups each male retrieved back to the nest, after they were removed from it, and the latent time it took for males to start retrieving them.

## METHODS

### Capture of wild mice

Mound-building mice were captured in northern Bulgaria in September 2011 and were then kept in the animal facilities of the University of Montpellier in France. We caught the mice in an agricultural area, at least 2 km away from the village of Rakita (GPS coordinates: 43°16'13.171”N 24°16'16.447”E), using Sherman live-traps. At this time of year, the mice had already built their mounds. Live-traps were set around each mound. A total of 30 juvenile males and 28 juvenile females captured from 14 mounds participated to this study. We weighed and measured all the mice from the nose to the base of tail upon capture. Same-mound mice, irrespective of their sex, were kept in the same laboratory cages from their capture until the start of the experiments two months later.

### Male cooperativeness during mound building

We used the contribution to mound building in large terrariums as a measure of male cooperativeness. This experiment lasted 8 weeks. Ten groups of three males and two females were constituted. Sex composition was based on the sex-ratio observed in the wild during the mound building period (Simeonovska-Nikolova and Gerasimov 2000; Simeonovska-Nikolova and Mehmed 2009). We placed each group in a large terrarium (1 × 1 × 0.8 m) containing approximately 0.2 m3 of earth and stones. The room was maintained at 21/23°C under a 12:12 light/dark cycle (corresponding roughly to the photoperiod observed in the field in September/November) with lights on at 9.00 pm. Water was available *ad libitum*, and food was provided on a weekly basis, although daily visits allowed checks on the well-being of the mice. At the beginning of each observation session, straw and seeds were provided as building materials in addition to the earth and stones already present in the terrariums. For each male, five categories of behavior were recorded: the number of times it dug, carried straw, carried seeds, carried stones, and entered the mound. Since the species is nocturnal, all the observations were conducted during the dark phase under dim red light. A single investigator (AT) observed each group during 5 sessions of 1 hour each, spaced out by about one week. Males were ear tagged (7.8 × 2.8 mm, Fine Science Tools GmbH). For each terrarium, the number of metallic earrings (1 or 2) and the side of the tag (right ear, left ear, or both) allowed individual mice to be recognized without having to handle them.

We could constitute only five groups of same-mound individuals (three males and two females). The other five groups included three same-mound males to limit aggression (Sokolov et al. 1998) and two same-mound females from a mound different from that of the males. Unfortunately, for four out of these five groups, we observed high female mortality (n = 8), most likely killed by the males during the first night in the enclosure. Following this observation, we immediately removed the two remaining females from the fifth terrarium and placed them in a separate cage.

We averaged the data collected during the five observation sessions for each male and for each of the behavioral items recorded during mound building (transport of straw, seeds, and stone, digging and entering into the mound). Because 22 out of 30 mice did not transport any seed, we removed this variable from the analysis. A principal component analysis (PCA) was applied to the four behavioral items to extract a single factor reflecting male propensity to invest in mound building. This factor was used as a measure of male individual cooperativeness.

At the end of this experiment, all individuals were transferred to laboratory cages, and females and males of a given group were kept in separate cages to prevent reproduction. They were maintained in laboratory conditions under a 13.5 h:10.5 h light/dark cycle (light at 6.30 pm), corresponding to the photoperiod during the breeding season (early spring). Food and water were available *ad libitum*, and cotton was provided as nesting material.

### Female preference

We measured female preference in a Y maze following the procedure described in Smadja & Ganem (2002; Smadja and Ganem 2005). Each female was tested once. The stimuli were composed of 5 pairs of males. Males of each pair shared the same terrarium during the mound building experiment and belonged to one of the five terrariums in which females were present. In each of the five terrariums, we selected among the three males the most and the least We could constitute only five groups of same-mound individuals (three males and two cooperative male in order to present a choice in cooperativeness to the females. To that aim, we used the factor reflecting male investment in mound building of the PCA (see above).

We tested the female preference for high over low cooperative males when cooperation was observed (the females shared the same enclosure with the male stimuli during mound building), and when cooperation was unobserved (the females were not present during mound building).

A single investigator (AT) conducted all the preference tests. All mice were more than 6 months old and sexually mature. To maximize the expression of sexual preferences, females were tested while sexually receptive (i.e., estrus or proestrus/estrus, assessed with vaginal smears).

The Y maze was transparent (plexiglas and plastic ware) and composed of a main branch (5 cm diameter, 35 cm long) connected to two secondary branches (5 cm diameter, 25 cm long). Boxes (35 × 23 × 13 cm) with transparent perforated doors were connected at the end of each branch [for an illustration of the apparatus see: 51]. One week before the experiment, each female was allowed to explore the empty Y maze for 15 minutes in order to become habituated.

At the start of each test, a female was placed in a box connected to the main branch of the apparatus. The two stimuli were randomly assigned to one or the other peripheral boxes (the identity of the stimuli was not known by the observer). We then opened the door of the female box and started to record its behavior when the female crossed the box door. In all tests, the females entered both secondary branches of the Y maze. During the 10-minute observation, we recorded the time spent by females: i) in each secondary branch (including when females were in contact with the perforated door in absence of a male just behind the door), and ii) sniffing or licking the transparent perforated door in the presence of a male just behind the door or interacting with the male. We used a paired Wilcoxon rank test (two-tailed tests, function *wilcox.test* in R) to test whether females spent more time on the side of the high versus low cooperative male. At the end of the experiment, all individuals were returned to their home cage and maintained under laboratory conditions.

### Measure of paternal investment through pup attendance during retrieval experiments

We used a pup retrieval procedure, a test used commonly to measure parental care in rodents (Dudley 1974; Cohen-Salmon et al. 1985), including *Mus spicilegus* (Patris and Baudoin 2000). We first randomly paired females (n = 20) and 'unfamiliar' males (i.e., captured in different mounds in the field). During the first week, the two partners were maintained in the same cage but separated with a wire net so that they could become familiar with each other. We maintained them in laboratory conditions under a 13.5:10.5 light/dark cycle (light at 6.30 pm), corresponding to the photoperiod of the breeding season in the field. Food and water were available *ad libitum*, and cotton was provided as nesting material. Ten out of the 20 couples successfully bred and could hence participate in this experiment. Litter size varied from 5 to 12 pups. We tested each male during two sessions with a two day interval.

On the day of birth, we placed the mice in a large terrarium (70 × 30 × 30 cm). The terrarium contained clean sawdust, food and water. One corner also contained cotton and cardboard rolls as nest-building material.

All males were tested twice: approximately four days (mean ± sd: 3.7 ± 0.8 days) and six days (6.3 ± 0.9 days) after birth of their first litter. During these two sessions, we first removed the breeding pair from the terrarium. We then removed three of the pups from the nest and placed them at the opposite end of the terrarium. We isolated these three pups from their littermates and the nest by placing a transparent plastic separation in the middle of the terrarium. The male was then put back in the side of the terrarium containing the nest. After 30 seconds, we removed the plastic separation and we measured (i) male latency before the start of retrieval of the isolated pups, and (ii) the number of pups retrieved to the nest during the next fifteen minutes. Both of these measures were averaged over the two test sessions, and we used Spearman correlation tests (two-tailed tests, function *cor.test* in R) to assess their potential association with the male cooperativeness score measured during the mound building experiment (first experiment above). For males who did not retrieve any pups, the male latency to retrieve was set at 15 minutes (i.e., the duration of the experiment). Because the Spearman correlation test relies on rank, this choice did not influence the results.

Because this species is particularly active during the first few dark hours (Simeonovska-Nikolova and Mehmed 2009), all observations (mound building, female preferences, and paternal investment) were conducted one to two hours after the beginning of the dark period. We recorded all observations using the Observer software Version 5 (Noldus Information Technology). All statistical analyses were performed using the R software, version 3.1.2 (R Core Team 2013).

### Ethical note

Mouse sampling was performed with the authorization of the Bulgarian Ministry of the Environment and Water (permit N°33-00-140), and housing and testing in Montpellier were authorized by the French authorities (permit N°C34-265). This study followed the ABS/ASAB guidelines for the ethical treatment of animals. We were particularly committed to limiting the number of mice trapped and tested, and mice were provided an enriched environment and diversified food to reduce their stress as much as possible.

## RESULTS

### Male cooperativeness

In total, each male transported 0 to 70 pieces of straw (mean ± s.d: 8.0 ± 19.4), and 1 to 99 (20.9 ± 23.9) stones (approximately 1 cm3 of size). The frequency of digging varied between 0 and 35 (10.1 ± 8.9), and they entered the mound 2 to 35 times (12.6 ± 9.9). The first two axes of a PCA, including the four mound building items, captured 80% of the total variation. The first axis was positively correlated with the number of times a male entered the mound and negatively correlated with the transport of stones and digging frequency (Table S1). The second axis was positively correlated with all four variables (frequency of digging, number of stones and straw transported, number of entrances into the mound) (Table S1). Hence, we considered that the second axis represented a measure of global investment in mound building, and thus, cooperativeness.

Cooperativeness was not related to male weight (10.26±0.87 grams) or size (7.14±0.34 cm) at capture (Spearman correlation test: weight: ρ = 0.11, *p* = 0.58, size: ρ = 0.01, *p* = 0.95) nor to their weight (11.26±1.6 grams) at the end of the mound building experiment (ρ = 0.13, *p* = 0.53). Moreover, we did not detect any difference in individual cooperativeness when comparing males (n = 15) belonging to mixed groups (containing females) to males belonging to all-male groups (n = 15) (Wilcoxon rank sum test: W = 86, *p* = 0.43).

### Female preference

Both females who observed male cooperation (n = 10) and those who did not (n = 10) spent significantly more time interacting with the high than with the low cooperative males through the perforated doors (paired Wilcoxon rank test: median of the difference [1st quartile; 3rd quartile] = 6.5 [0.17; 9.4] s, V = 40, *p* = 0.04; 4.6 [0.0; 10.5] s, V = 21, *p* = 0.04, respectively; Fig. 1A). However, they explored the two branches leading to the male cages to the same extent (for females who observed male cooperation or not, respectively: 4.2 [-6.4; 18.2] s, V = 35, *p* = 0.49; 18.2 [- 9.0; 104.7] s, V = 36, *p* = 0.43; Fig. 1B).

**Figure 1.**
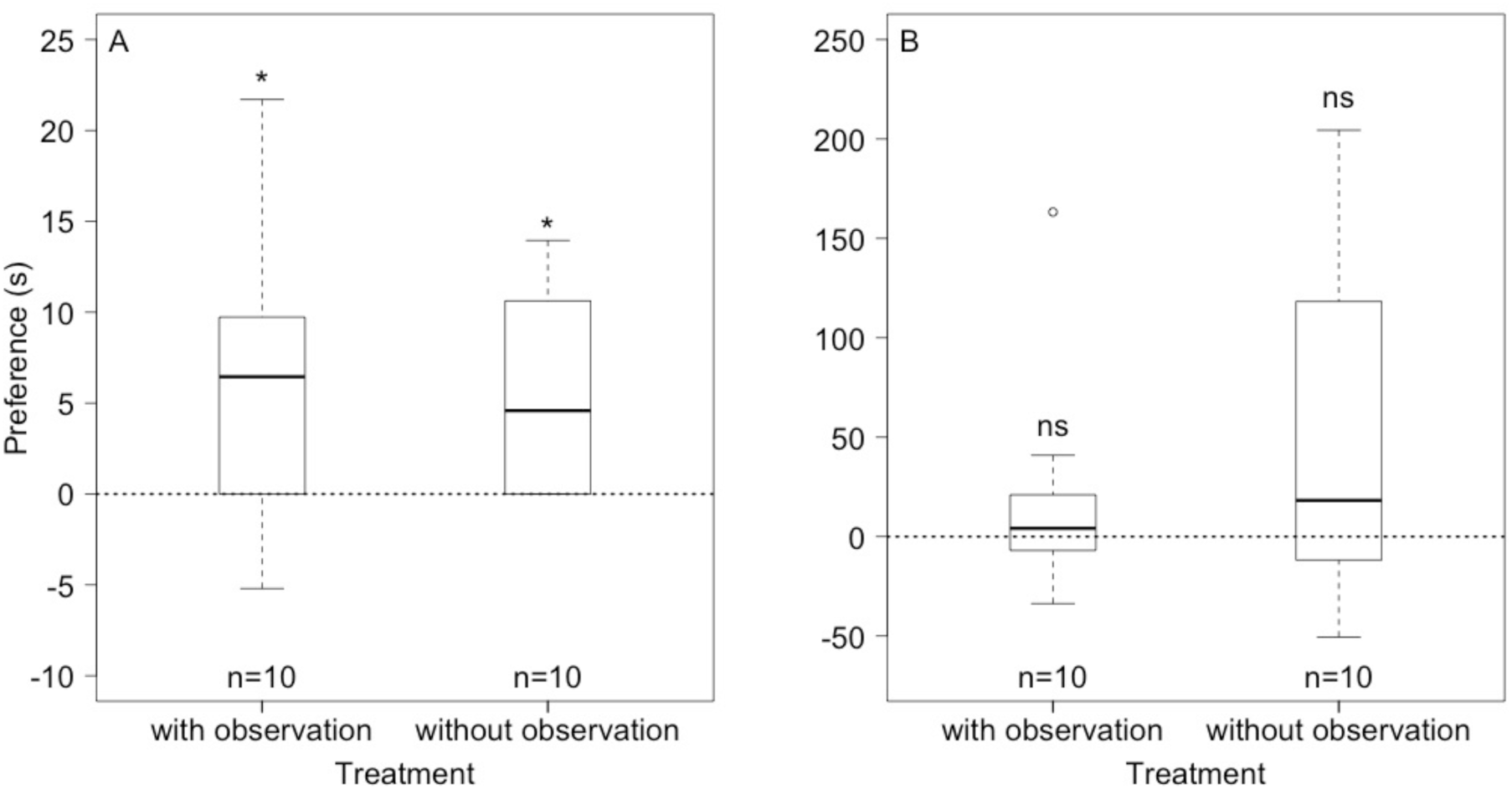
Preference of females who did (n = 10) or did not (n = 10) observe male mound building. Preference is calculated as the difference in time (in seconds) spent interacting through the perforated door with the high versus the low cooperative males (A) and the difference in time spent in the branch leading to the box of the strongly versus weakly cooperative males (B). When the cooperative male is preferred, the difference is positive. The horizontal dotted line represents random choice (preference = 0). Box-plots show the preference median (thick line) and first and third quartiles. ns: non-significant, * p<0.05.

### Cooperativeness & paternal investment

The males (n = 10) retrieved 0 to 3 pups for each session (median [1st quartile; 3rd quartile]: 1.5 [0; 3] for the first session; 2.5 [0.25; 3] for the second session). For each male, the number of pups retrieved during the first versus the second session was not significantly different (paired Wilcoxon rank test: 0 [-0.75; 0] pups, V = 5.5, *p* = 0.68). For sessions in which at least one pup was retrieved (n = 12), the latency to retrieve the first pup was between 23 and 810 seconds (171 [36; 325] s). Contrary to our prediction, paternal attendance was negatively associated with male cooperativeness: across the 20 sessions, males (n = 10) investing more in mound building retrieved fewer pups back to the nest (Spearman correlation test: ρ =-0.67, *p* = 0.04) and presented a higher latency time to retrieve the first pup (ρ = 0.71, *p* = 0.02).

## Discussion

Although indirect fitness benefits largely explain the evolution of cooperation in a broad range of species (Clutton-Brock 2002; Griffin and West 2002; West et al. 2007; Bourke 2014; Dijk et al. 2014), other mechanisms, such as mate preference, can help us understand how non-kin cooperative interactions may evolve (Putland 2001; Nowak 2006; Covas et al. 2007; Nesse 2007; McNamara et al. 2008; Barclay 2011). Empirical studies in humans and observations in some cooperative birds suggest that cooperative behaviors may act as signals during mate choice (Reyer 1984; Jones 1998; Doutrelant and Covas 2007; Tognetti et al. 2012; Van Vugt and Iredale 2013; Raihani and Smith 2015; Tognetti et al. 2016). However, cooperative behaviors are not always observable by potential partners before mate choice. Using two-way choice tests, we investigated whether females preferred strongly over weakly cooperative males in the mound-building mouse even though females did not observe male mound-building behaviors. The overwinter survival of offspring relies on mound protection, and we also assessed whether highly cooperative males were more involved in offspring attendance using pup-retrieval experiments. Our findings show that females prefer high over low cooperative males, even when cooperative behaviors had not been observed. This suggests that female mate choice is influenced by cues of cooperativeness that are different than cooperative behaviors *per se*, or by preferences for traits associated with male cooperativeness. We also show that male attendance to offspring is negatively correlated with male cooperativeness, suggesting the existence of alternative strategies of offspring care.

Our findings raise the question of which signals or cues females are responding to when they show a preference for cooperative males. In our experiment, females are more attracted to high over low cooperative males, even without having observed them build. Hence, phenotypic cues advertising male cooperativeness may exist in this species, as previously found in humans (Oda et al. 2009; Stirrat and Perrett 2010; Stirrat and Perrett 2012; Tognetti et al. 2013; Centorrino et al. 2015) and in barn owls (Roulin et al. 2012). Such cues of cooperativeness could be based on acoustic, olfactory, or visual traits, since all of them are involved in mice communication (e.g., Hurst and Beynon 2004; Musolf et al. 2010) and are available to females in our apparatus. Their detection by females could trigger an innate response, similarly to responses to some sexual pheromones (Roberts et al. 2010; Li and Liberles) or predators' cues (Veen et al. 2000; Hawkins et al. 2004; Fendt 2006; Ferrero et al. 2011) observed in several species, including mice. An alternative explanation could also be that female preference for strongly over weakly cooperative males was not directed to cooperativeness *per se* but to other traits with which cooperativeness is associated. We did not observe any link between male size or weight and cooperativeness, suggesting that male condition is unlikely to be one of these traits. Interestingly, experimental studies in the mound-building mouse found that agonistic behaviors between unfamiliar males and females are positively linked with sexual motivation (Busquet et al. 2009; Simeonovska-Nikolova and Lomlieva 2012). Because of the probable high cost of mound building and its important role in the success of overwintering, individuals who invest more in mound building are likely to more aggressively protect resources and defend their mound from intruders. Additionally, dominant males could also invest more in mound building because they might get more benefits. We can therefore speculate that male aggressiveness or dominance might be sexually selected traits associated with cooperativeness, leading to the indirect selection of cooperativeness through female mate choice. During our preference tests, we did not observe agonistic behaviors between males and females through the perforated doors. However, chemosensory cues of dominance and aggressiveness are present in the urine and preputial glands of male mice and therefore could have been detectable by females during their interactions with the males (Harvey et al. 1989; Hurst and Beynon 2004; Mucignat-Caretta and Caretta 2014). Follow-up studies should thus investigate which traits, among those used by females *Mus spicilegus* during mate choice, could be associated with cooperativeness. Such studies may particularly focus on dominance, aggressiveness and other personality traits.

In *Mus spicilegus*, the inclination of females to prefer cooperative males could be an adaptive strategy. Choosing a cooperative male could increase females' fitness in several ways. First, since mound size is positively correlated to water-insulation and soil temperature inside the mound (Szenczi et al. 2011; Szenczi et al. 2012), choosing a male who invests more energy and time in mound building could increase offspring survival during the winter. Second, mound building behavior seems to be genetically heritable (Orsini 1982). By choosing a cooperative male, females could improve the ability of their offspring to build a mound during the following summer, and hence, increase the chance of survival for their own progeny. Third, given the attractiveness of highly cooperative males, females could also increase their sons' reproductive success (sexy son hypothesis: Weatherhead and Robertson 1979).

Interestingly, our results note a negative correlation between offspring attendance and cooperativeness. Previous experimental studies involving the mound building mouse found intense paternal investment, such as covering and warming the pups (Patris and Baudoin 2000), and males that exhibited the highest levels of offspring attendance increased their reproductive success by reducing their mate's inter-litter intervals (Feron and Gouat 2007). As both parental investment and cooperative behavior in mound building are likely to be energetically costly, the negative association observed between them may indicate the existence of a tradeoff between two alternative parental care strategies. Hence, an interpretation of our results could be that different behavioral traits may correspond to different types of fathers with regard to how they take care of offspring. Such alternative strategies in offspring care were previously observed in some cooperative breeding species, such as the cichlid, *Neolamprologus pulcher*, or the noisy miner, *Manorina melanocephala*, in which different types of helpers seem to exist (Arnold et al. 2005; Bergmüller and Taborsky 2007), but to our knowledge, they were not extensively studied in relation to parental care.

Overall, our results provide rare support for the hypothesis that cooperativeness could be detectable by phenotypic cues different from cooperative behaviors *per se* and that these cues can influence mate choice. While the mechanisms enabling females to prefer cooperative males in this species are still unknown, our results suggest that mate choice might influence the evolution and maintenance of cooperative behavior and that sexual selection may be another pathway by which cooperation evolved.

## Acknowledgments

We are very grateful to D. Simeonovska-Nikolova, B. Nikolovi and C. Albinet for their help for the trapping session in Bulgaria. We also thank Z. Groó, Y. Latour, A. Orth, and M. Perriat-Sanguinet, J. Barthes, J. Bovet, M. Derex, L. Danilo, V. Durand, L. Etienne, A. Nitsch, and A. Ravel, to help provided throughout the experiment. We thank the Ministry of Environment and Water of Bulgaria, which permitted us to catch specimen of *Mus spicilegus*. Supports through the ANR-Labex IAST, the CNRS of France (www.cnrs.fr), the Région Languedoc-Roussillon 'Chercheur(se)s d'Avenir' (no.: DGA3/DESR/2012/Q159) and the Fondation des Treilles (www.lestreilles.com) are gratefully acknowledged.

